# Genetic legacy in soil seedbanks after grassland conversion to plantation forests: evidence from *Potentilla freyniana*

**DOI:** 10.64898/2026.08.28.747765

**Authors:** Suzuki Setsuko, Kentaro Uchiyama, Tomoyo Koyanagi, Kei Uchida, Fujio Hyodo, Fangzheng Fu, Asuka Koyama

## Abstract

Semi-natural grasslands are important ecosystems supporting biodiversity in Japan, but their area has declined rapidly due to land-use change and abandonment of traditional management practices such as mowing and burning. Although the conservation of genetic diversity is essential for the long-term persistence of grassland plants, little is known about the genetic diversity retained in soil seedbanks following conversion of grasslands to plantation forests. In this study, we compared the genetic diversity and population structure of above-ground and soil seedbank populations of the grassland perennial forb *Potentilla freyniana* across three sites in each of three land-use types: burned grasslands, deciduous plantation forests, and evergreen plantation forests (plantation ages approximately 21–62 years) on the Kaida Plateau, central Japan. Soil seedbank populations were obtained from soil samples through germination experiments, and genetic analyses were conducted using newly developed simple sequence repeat (SSR) markers. Genetic diversity was assessed using expected heterozygosity, allelic richness, and private allelic richness, population structure was evaluated using analysis of molecular variance (AMOVA), STRUCTURE analyses, and pairwise *F*_ST_. Soil seedbank populations maintained levels of genetic diversity comparable to those of above-ground populations, and no significant differences were detected between the two population types. Furthermore, soil seedbank populations in evergreen plantation forests, where above-ground individuals of *P. freyniana* were absent, retained genetic diversity comparable to that observed in burned grasslands. AMOVA detected no significant genetic differentiation between above-ground and soil seedbank populations. These results suggest that high levels of genetic diversity can persist in soil seedbank populations for decades after forest establishment and highlight the potential importance of soil seedbanks as genetic resources for grassland restoration.

## Introduction

Soil seedbanks, defined as seed assemblages stored in a dormant state within the soil, constitute an important ecological component that supports the persistence of plant populations and vegetation recovery. Soil seedbanks are widely recognized as a temporal bet-hedging strategy against environmental uncertainty and are thought to contribute to the long-term persistence of species [1]. In addition, soil seedbanks may increase effective population size and help maintain genetic diversity by buffering populations against genetic drift and demographic fluctuations [2]. They also serve as important reservoirs of plant diversity and can strongly influence future vegetation composition and ecosystem recovery [3]. Moreover, soil seedbanks can function as a genetic memory that preserves historical genetic diversity, often retaining levels of diversity comparable to those of above-ground populations while exhibiting differences in allele frequencies [4]. Together, these studies indicate that soil seedbanks are important components that integrate both contemporary population dynamics and historical genetic processes.

Semi-natural grasslands, on the other hand, are ecosystems that have been maintained through traditional land-use practices across many regions of the world and provide important habitats for diverse grassland plants and insect communities [5]. However, many semi-natural grasslands have declined because of land-use change and management abandonment worldwide [6]. In Japan, semi-natural grasslands are secondary ecosystems maintained through anthropogenic management practices such as controlled burning, mowing, and grazing, and they have long served as important habitats for a wide variety of grassland plant species. Although semi-natural grasslands covering substantial areas were historically maintained in mountainous regions of Japan, their extent has declined dramatically over the past century because of afforestation and natural forest expansion following management abandonment [7]. Consequently, many former grasslands are now occupied by plantation forests, raising the question of whether grassland biodiversity and genetic resources persist in soil seedbanks in these transformed habitats.

Ecological studies have highlighted the importance of legacy effects, whereby past land-use history exerts long-lasting influences on vegetation structure and species diversity [8, 9]. In addition, attention has been paid not only to above-ground vegetation but also to soil seedbanks below ground, which play a crucial role in ecosystem resilience and recovery [10, 11]. Previous studies have shown that soil seedbanks may serve as important sources for grassland recovery following afforestation. However, the contribution of soil seedbanks to grassland restoration is often limited because seeds of grassland species gradually disappear from the soil after afforestation [12–14]. Further studies at both the community and species levels are needed to clarify the contribution of soil seedbanks to grassland restoration, particularly in areas where populations of target grassland species have disappeared from the above-ground vegetation.

Nevertheless, most studies on soil seedbanks have focused on ecological attributes such as species composition and diversity [10], whereas relatively few studies have examined the genetic diversity maintained within soil seedbanks and their genetic relationships with extant above-ground populations [15–17]. Although persistent soil seedbanks have been suggested to mitigate the effects of habitat fragmentation by reducing genetic drift and population genetic differentiation [18], few studies have evaluated genetic diversity in both soil seedbank and above-ground populations across different land-use conditions, particularly following the conversion of grasslands to plantation forests. Consequently, our understanding remains limited regarding the extent to which soil seedbanks retain genetic variation following the conversion of grasslands to forest plantations and how this genetic variation relates to present above-ground populations.

The Kaida Plateau in Nagano Prefecture, Japan was once characterized by extensive semi-natural grasslands maintained through traditional practices such as Kiso horse grazing and hay harvesting. However, grassland area declined dramatically from approximately 5,000 ha in 1955 to only about 5 ha in 2021 as a result of land-use change [19]. Under such drastic landscape transformation, it remains unclear how grassland plant populations have been genetically affected and whether soil seedbanks persisting beneath forest plantation sites still retain historical genetic diversity.

In this study, we investigated *Potentilla freyniana* Bornm. (Rosaceae), a characteristic perennial grassland species occurring in Japanese semi-natural grasslands and compared the genetic diversity and genetic structure of above-ground populations and soil seedbank populations. Previous studies have shown that *P. freyniana* can be detected in soil seedbanks even after populations have disappeared from the above-ground vegetation, suggesting that this species can form a long-term persistent seedbank [20, 21]. Specifically, we aimed to determine whether soil seedbanks in forest plantation sites maintain high levels of genetic diversity and to what extent they remain genetically continuous with contemporary above-ground populations. By doing so, we evaluated the persistence of genetic legacy following land-use change and assessed the potential value of soil seedbanks for grassland conservation and restoration.

## Materials and Methods

### 1. Study sites

The study was conducted in the Kaida Plateau, Nagano Prefecture, central Japan, at elevations ranging from 1,140 to 1,250 m a.s.l. The landscape consists of a forest-grassland mosaic that historically supported extensive semi-natural grasslands. Analysis of aerial photographs revealed that semi-natural grasslands covered approximately 20 % of the study area in 1948, whereas forests occupied 68 % of the landscape. By 2010, semi-natural grasslands had declined to approximately 2 % of the study area, largely as a result of conversion to plantation forests [22]. Consequently, the remaining grasslands are small, isolated, and distributed within a forest-dominated landscape. The semi-natural grasslands are dominated by *Miscanthus sinensis* and various perennial herbaceous species [23].

Study sites were established only in locations that were identified as semi-natural grasslands in the 1948 aerial photographs. All study sites were small, fragmented, and spatially isolated from one another (Fig. 1). Based on current land use, sites were classified into three categories: burned grassland (Bg), deciduous plantation (Dp; plantations of *Larix kaempferi*), and evergreen plantation (Ep; plantations of *Cryptomeria japonica* or *Chamaecyparis obtusa*). Burned grasslands have been continuously maintained through prescribed burning and, in some cases, mowing. Deciduous plantations consisted of monocultures of *L. kaempferi*, whereas evergreen plantations consisted of monocultures of *C. japonica* or *C. obtusa*. Plantation age ranged from approximately 21 to 62 years, and tree height ranged from approximately 15 to 30 m. Three sites were selected for each land-use category (Table 1, Fig. 1). We attempted to include all three land-use categories within the same locality whenever possible to reduce environmental heterogeneity among sites.

**Fig. 1.**
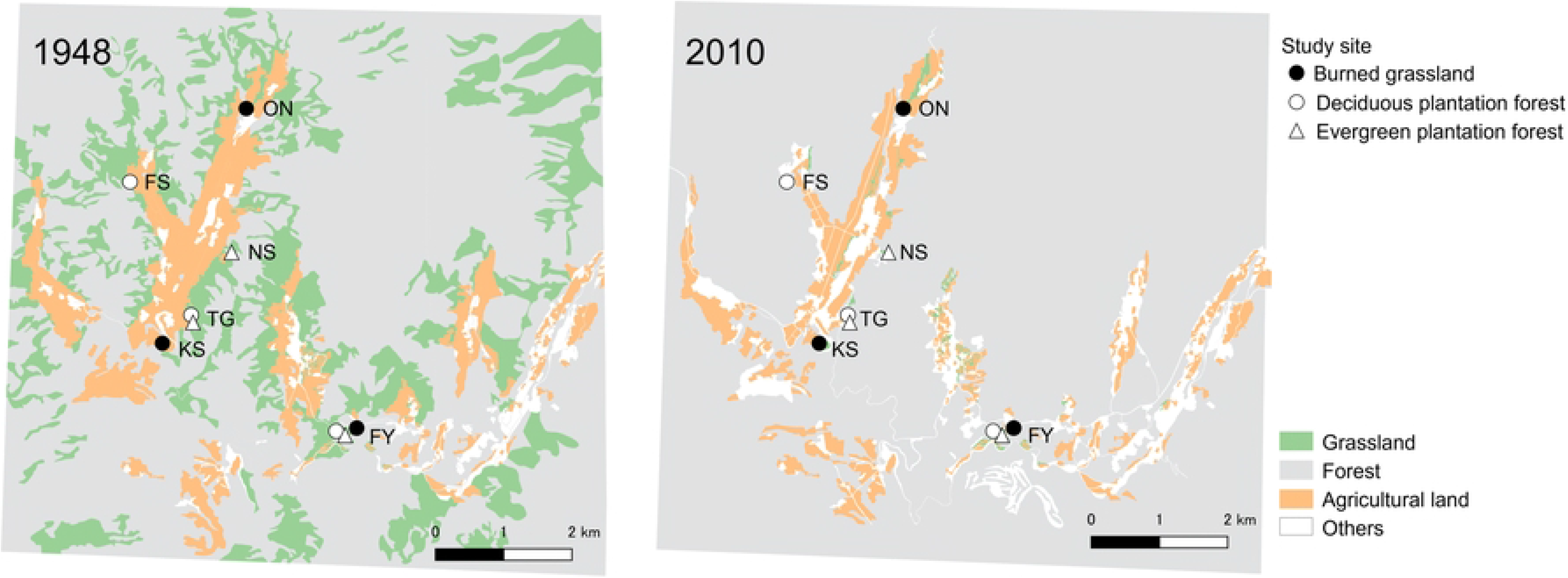
Land-use changes in the study area at Kaida Plateau, Nagano Prefecture, Japan, between 1948 and 2010 based on aerial photographs. Locations of the six study sites across three land-use types (Bg: burned grassland; Dp: deciduous plantation forest; Ep: evergreen plantation forest) are shown. All sites were managed grasslands in 1948. Adapted from Koyama and Uchida [11] and based on the land-use classification of Yamamoto and Uchida [22]. Base map data were provided by the Geospatial Information Authority of Japan (GSI).

**Table 1.**
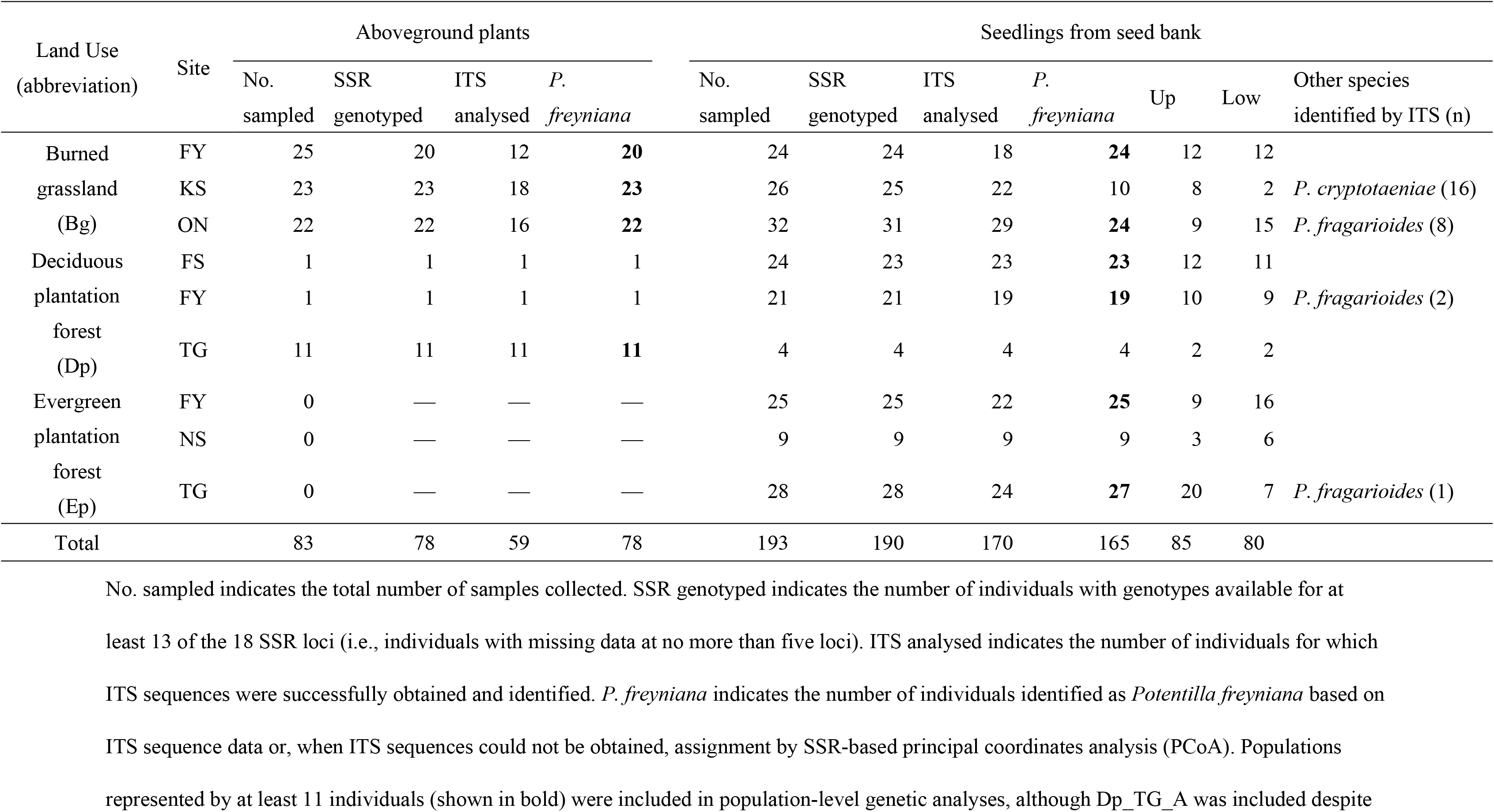
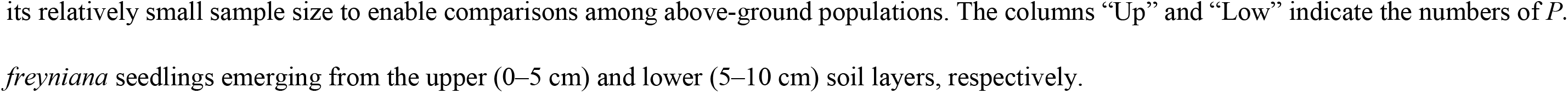
Sample sizes, genotyping success, and species identification of above-ground and seed-bank samples across land-use types and sites, with additional information on *Potentilla* species in the Kaida Plateau. No. sampled indicates the total number of samples collected. SSR genotyped indicates the number of individuals with genotypes available for at least 13 of the 18 SSR loci (i.e., individuals with missing data at no more than five loci). ITS analysed indicates the number of individuals for which ITS sequences were successfully obtained and identified. *P. freyniana* indicates the number of individuals identified as *Potentilla freyniana* based on ITS sequence data or, when ITS sequences could not be obtained, assignment by SSR-based principal coordinates analysis (PCoA). Populations represented by at least 11 individuals (shown in bold) were included in population-level genetic analyses, although Dp_TG_A was included despite its relatively small sample size to enable comparisons among above-ground populations. The columns “Up” and “Low” indicate the numbers of *P. freyniana* seedlings emerging from the upper (0–5 cm) and lower (5–10 cm) soil layers, respectively.

| Land Use<br>(abbreviation) | Site | Aboveground plants |  |  |  | Seedlings from seed bank |  |  |  |  |  |  |
| --- | --- | --- | --- | --- | --- | --- | --- | --- | --- | --- | --- | --- |
|  |  | No.<br>sampled | SSR<br>genotyped | ITS<br>analysed | *P.*<br>*freyniana* | No.<br>sampled | SSR<br>genotyped | ITS<br>analysed | *P.*<br>*freyniana* | Up | Low | Other species<br>identified by ITS (n) |
| Burned | FY | 25 | 20 | 12 | **20** | 24 | 24 | 18 | **24** | 12 | 12 |  |
| grassland | KS | 23 | 23 | 18 | **23** | 26 | 25 | 22 | 10 | 8 | 2 | *P. cryptotaeniae* (16) |
| (Bg) | ON | 22 | 22 | 16 | **22** | 32 | 31 | 29 | **24** | 9 | 15 | *P. fragarioides* (8) |
| Deciduous | FS | 1 | 1 | 1 | 1 | 24 | 23 | 23 | **23** | 12 | 11 |  |
| plantation | FY | 1 | 1 | 1 | 1 | 21 | 21 | 19 | **19** | 10 | 9 | *P. fragarioides* (2) |
| forest | TG | 11 | 11 | 11 | **11** | 4 | 4 | 4 | 4 | 2 | 2 |  |
| (Dp) |  |  |  |  |  |  |  |  |  |  |  |  |
| Evergreen | FY | 0 | — | — | — | 25 | 25 | 22 | **25** | 9 | 16 |  |
| plantation | NS | 0 | — | — | — | 9 | 9 | 9 | 9 | 3 | 6 |  |
| forest | TG | 0 | — | — | — | 28 | 28 | 24 | **27** | 20 | 7 | *P. fragarioides* (1) |
| (Ep) |  |  |  |  |  |  |  |  |  |  |  |  |
| Total |  | 83 | 78 | 59 | 78 | 193 | 190 | 170 | 165 | 85 | 80 |  |

However, all three categories were available within a single locality only at the FY site.

### 2. Sampling and germination experiment

*Potentilla freyniana* is a stoloniferous perennial forb distributed throughout East Asia, including Japan. It commonly occurs in semi-natural grasslands and open forests and is regarded as a characteristic grassland species. The species produces flowering stems 5–20 cm in height [24], and has small, lightweight seeds (mean seed mass: 0.22 mg) that are primarily dispersed by gravity [25], although the presence of elaiosomes suggests that ants may also contribute to seed dispersal [26]. The flowering period at the study area is from April to June.

Above-ground individuals of *P. freyniana* were sampled from all nine study sites in late June 2022. In populations with sufficient numbers of individuals, samples were collected across the population area while maintaining a minimum distance of approximately 2–3 m between sampled individuals to minimize spatial bias and reduce the likelihood of repeatedly sampling clonally connected ramets. Because population sizes of *P. freyniana* were small in the deciduous plantation sites, nearly all available individuals were sampled. To increase sample size, additional individuals were collected from the Dp_TG_A population in late June 2024 and combined with the 2022 samples for subsequent analyses. In total, leaf samples were collected from 22–25 individuals at each of the three burned-grassland sites and from 1–11 individuals at each of the three deciduous plantation sites (Table 1). No above-ground individuals were found in the evergreen plantation sites. In addition, leaf samples were collected from seven above-ground individuals of the closely related species *Potentilla fragarioides* occurring near the study area and used as a reference group for species identification. All leaf samples were transported to the Forestry and Forest Products Research Institute under cool conditions and stored frozen until DNA extraction.

To sample soil seedbank populations, soil samples were collected in late October 2022 and 2024 from the three land-use types described above. At each site, soil was collected from 20 sampling points located at least 5 m apart to maximize spatial coverage. Using a 100 cm^3^ soil core, soil samples were collected from two depth intervals (0–5 cm and 5–10 cm). In total, 18 L of soil was collected in each sampling year, resulting in 36 L across both years. Additional sampling was conducted in the second year because insufficient numbers of seedlings emerged from some sites during the first sampling year. For subsequent genetic analyses, seedlings emerging from the two soil depths were pooled and treated as a single soil seedbank population in order to obtain sufficient sample sizes for population genetic analyses. This approach was supported by the absence of significant differences in expected heterozygosity (*H*_E_) between the upper and lower soil layers at Bg_FY_S and Dp_FS_S where sufficient numbers of seedlings were available from both depths (Table 1).

The soil samples were stored in plastic bags and cold-stratified under moist conditions at 3 °C in a dark incubator. In February of the year following collection, soil samples were processed for germination experiments in a greenhouse at the Forestry and Forest Products Research Institute. Prior to sowing, soil samples were passed through a 2- mm sieve to remove stones and separate vegetative propagules from buried seeds. The processed soil was then spread in trays and placed in a greenhouse covered with white mesh netting to prevent contamination by external seeds. The soil was watered regularly throughout the experiment.

Leaf tissue was collected from emerged seedlings, with a maximum of four individuals from each soil sample. As a result, 24–32 seedlings were obtained from the three burned-grassland sites, 4–24 seedlings from the three deciduous plantation sites, and 9–28 seedlings from the three evergreen plantation sites (Table 1). Leaf tissue was harvested after the development of several true leaves and stored frozen until DNA extraction.

### 3. DNA extraction and species identification using the ITS1 region

Genomic DNA was extracted from frozen leaf tissue using a modified CTAB protocol [27]. Because leaves of *Potentilla* species contain high concentrations of polyphenols, tannins, and polysaccharides, multiple washing steps were required prior to DNA extraction with CTAB buffer. Species identification of individuals originating from the soil seedbank was primarily based on seedling morphology. However, discrimination among closely related species, such as *P. fragarioides*, can be difficult at the seedling stage. Therefore, species identification was supplemented using sequence information from the internal transcribed spacer (ITS) region. Although the genus *Potentilla* includes numerous species, most can be readily distinguished because their ITS sequences show clear interspecific differences. In contrast, *P. freyniana* and *P. fragarioides* exhibit highly similar ITS sequences, making reliable discrimination more challenging. Accordingly, we focused on subtle sequence differences between these two species and identified diagnostic nucleotide substitutions useful for species identification.

ITS sequences were obtained from *P. freyniana* individuals collected in Nagano, Ibaraki, and Tokyo Prefectures, and from *P. fragarioides* individuals collected in Nagano and Shizuoka Prefectures. These sequences were further compared with homologous sequences of overseas origin available in the NCBI database. As a result, two diagnostic nucleotide substitutions were identified within the ITS1 region that consistently distinguished the two species (Table S1), and these species-specific differences were conserved across geographically distant samples. However, after increasing the number of samples examined, a small number of *P. freyniana* individuals heterozygous at position 114 of the ITS1 region were detected, indicating that this site was not a reliable diagnostic marker. In contrast, the nucleotide substitution at position 127 of the ITS1 region consistently differentiated the two species across all samples examined. Therefore, this position was used as the diagnostic nucleotide substitution for species identification in this study.

ITS1 amplification and sequencing were attempted for all above-ground individuals and soil seedbank-derived individuals. Species identification was performed by comparing the obtained sequences with reference sequences. In cases where a sequence did not clearly match either *P. freyniana* or *P. fragarioides*, a BLASTn search was conducted, and the species exhibiting the highest sequence similarity was assigned as the identification result. For individuals from which ITS1 sequences could not be obtained due to PCR amplification failure or other technical reasons, principal coordinates analysis (PCoA) based on simple sequence repeat (SSR) markers (described below) was conducted. Individuals clustering with samples that had been confirmed as *P. freyniana* based on ITS sequences were treated as *P. freyniana* and included in subsequent analyses.

### 4. SSR marker development

To develop polymorphic SSR markers for *P. freyniana*, next-generation sequencing was performed using genomic DNA extracted from a *P. freyniana* individual collected in the Kaida Plateau. Library preparation and sequencing were outsourced to Novogene Co., Ltd. Genomic libraries were constructed using the NEBNext Ultra II DNA Library Prep Kit (New England Biolabs) from DNA fragments with an average insert size of approximately 300 bp. Sequencing was conducted on an Illumina NovaSeq 6000 platform using 150-bp paired-end reads (PE150).

Raw sequence reads were processed using CLC Genomics Workbench ver. 12 (QIAGEN). After removing adapter sequences and low-quality regions, overlapping paired-end reads were merged. The merged and unmerged reads were then combined and used for subsequent analyses. Identification of SSR loci and primer design were conducted using QDD [28] with default settings. Of the 13,326,488 reads analyzed, SSR motifs were detected in 427,862 reads, yielding 58,846 unique sequences. Primer design using QDD generated 28,308 candidate SSR primer pairs.

From these candidates, 96 primer pairs were selected based on QDD design category, alignment score, number of SSR repeats, and predicted PCR product size. Following PCR screening, 18 loci showing consistent amplification and sufficient polymorphism were retained for subsequent genetic analyses (Table 2).

**Table 2.**
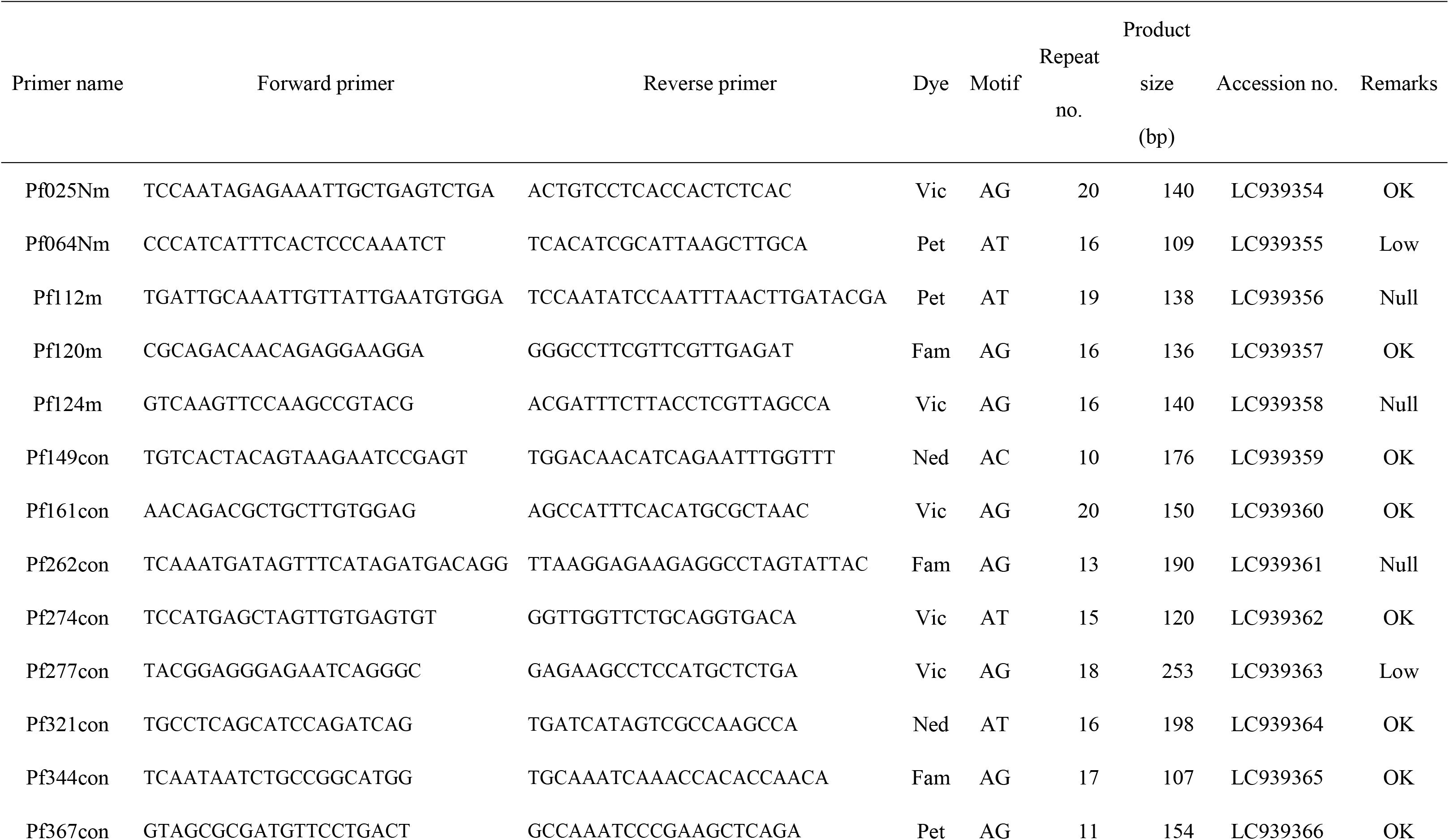
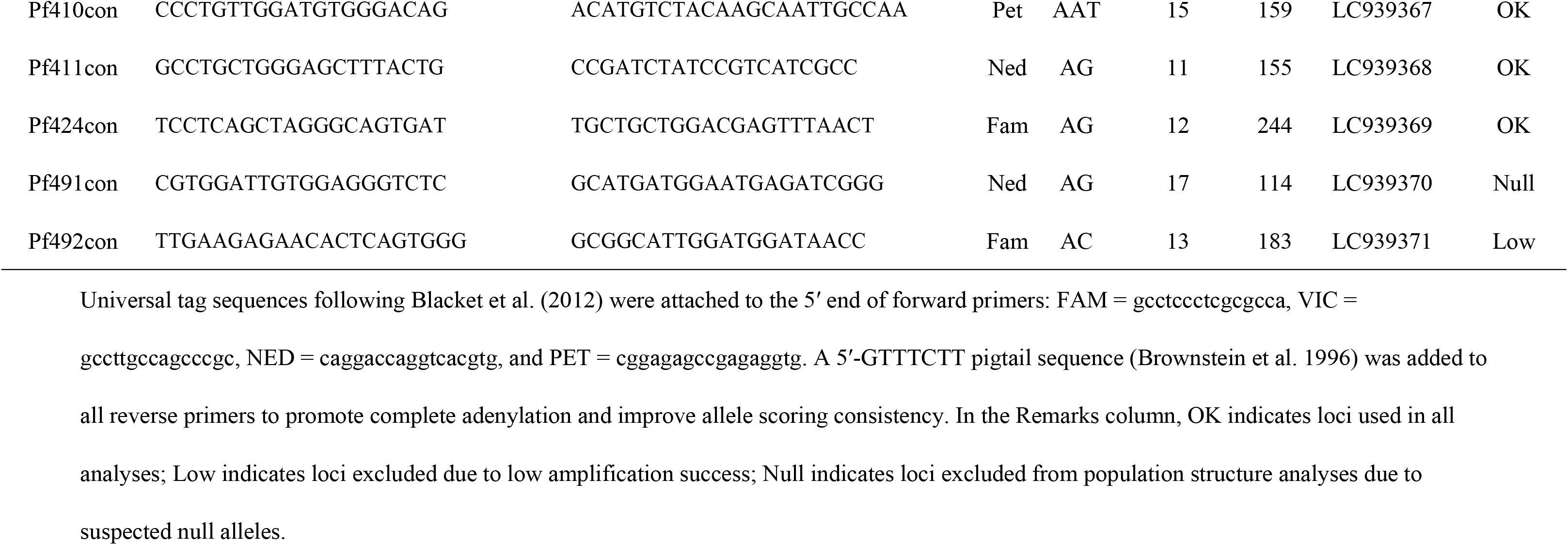
Characteristics of newly developed simple sequence repeat (SSR) markers for *Potentilla freyniana*, including primer sequences, repeat motifs, and amplification properties. Universal tag sequences following Blacket et al. (2012) were attached to the 5′ end of forward primers: FAM = gcctccctcgcgcca, VIC = gccttgccagcccgc, NED = caggaccaggtcacgtg, and PET = cggagagccgagaggtg. A 5′-GTTTCTT pigtail sequence (Brownstein et al. 1996) was added to all reverse primers to promote complete adenylation and improve allele scoring consistency. In the Remarks column, OK indicates loci used in all analyses; Low indicates loci excluded due to low amplification success; Null indicates loci excluded from population structure analyses due to suspected null alleles.

### 5. SSR analysis

PCR amplification was performed using the 18 selected SSR loci (Table 2). PCR products were separated using an ABI PRISM 3130 Genetic Analyzer (Applied Biosystems), and genotypes were scored using GeneMarker software (SoftGenetics).

The resulting genotype dataset was examined for missing data. Three loci with high proportions of missing genotypes were excluded, and the remaining 15 loci were retained for analyses of genetic diversity. In addition, inbreeding coefficients (*F*_IS_) were calculated for each population using FSTAT ver. 2.9.4 [29, 30]. Four loci exhibiting substantial heterozygote deficiency in more than half of the populations were suspected to contain null alleles and were therefore excluded from analyses of population structure to avoid potential bias.

Consequently, all genetic diversity analyses, including expected heterozygosity (*H*_E_), allelic richness (*A*_R_), and private allelic richness (PAR), were conducted using 15 loci. In contrast, population structure analyses, including PCoA, analysis of molecular variance (AMOVA), pairwise fixation indices (*F*_ST_) estimation, STRUCTURE analysis, and calculation of Jaccard coefficients, were performed using the remaining 11 loci.

### 6. Genetic diversity analyses

Genetic diversity within each population was quantified using *H*_E_, calculated in GenAlEx ver. 6.5 [31]. *A*_R_ and PAR were estimated using HP-RARE [32], with sample-size differences corrected by rarefaction. To ensure reliable estimates of genetic diversity, analyses were generally restricted to populations represented by at least 15 individuals. However, the above-ground population from deciduous plantations (Dp_TG_A), which consisted of 11 individuals, was retained because it provided an important comparison with above-ground populations in burned grasslands.

Differences in *H*_E_ and *A*_R_ among populations were first evaluated. Subsequently, populations were grouped according to land-use type (burned grassland, deciduous plantation, and evergreen plantation) and sample type (above-ground population and soil seedbank population), and *H*_E_ and *A*_R_ were compared among these categories.

Differences in genetic diversity among populations and categories were tested using linear mixed models (LMMs). All analyses were conducted in R version 4.3.2 [33] using the lme4 package [34]. Population or category was included as a fixed effect, whereas locus was included as a random effect to account for repeated measurements across loci.

The significance of fixed effects was assessed by analysis of variance (ANOVA) using the lmerTest package [35]. When significant differences were detected, post hoc multiple comparisons were conducted using Tukey’s method implemented in the emmeans package [36].

Because PAR is strongly influenced by geographic variation, comparisons were restricted to paired above-ground and soil seedbank populations within the same locality. Specifically, analyses were conducted using the FY site, where all three land-use types were represented, and the ON burned-grassland site, where sufficient numbers of both above- ground and soil seedbank samples were obtained. PAR values estimated by HP-RARE were compared using paired Wilcoxon signed-rank tests, treating individual SSR loci as replicates. Statistical significance was assessed at α = 0.05.

### 7. Population structure analyses

Genetic differentiation among populations was evaluated by calculating pairwise *F*_ST_ values using FSTAT ver. 2.9.4. Statistical significance was assessed using permutation tests.

To examine the hierarchical partitioning of genetic variation, AMOVA was conducted in GenAlEx ver. 6.5. Because the primary objective of this study was to evaluate genetic continuity between above-ground and soil seedbank populations, sample type (above-ground population vs. soil seedbank population) was used as the highest hierarchical level. AMOVA was performed using the 11 loci retained after excluding loci suspected of containing null alleles. Genetic variation was partitioned among sample types (above- ground populations and soil seedbank populations), among sampling sites within sample types, and among individuals within sites.

Population structure was inferred using STRUCTURE ver. 2.3.4 [37]. Analyses were conducted under the admixture model with correlated allele frequencies. The number of genetic clusters (*K*) varied from 1 to 12, and ten independent runs were performed for each value of *K*. Each run consisted of a burn-in period of 50,000 iterations followed by 100,000 Markov chain Monte Carlo (MCMC) iterations. The optimal number of clusters was determined using the Δ *K* method of Evanno et al. [38]. Results from replicate runs were summarized and visualized using CLUMPAK [39].

To evaluate allelic similarity among above-ground and soil seedbank populations, Jaccard coefficients were calculated based on the presence or absence of alleles detected in each population. For each allele, populations in which the allele was observed were assigned a value of 1, whereas populations lacking the allele were assigned a value of 0. Similarity between pairs of populations was then calculated as the number of shared alleles divided by the total number of alleles present in either population. Jaccard coefficients were visualized as heatmaps representing shared-allele similarity. All analyses were conducted in R version 4.3.2. Because the above-ground population in deciduous plantations (Dp_TG_A) consisted of relatively few individuals, and allele detection may therefore have been incomplete, this population was excluded from the Jaccard similarity analysis.

## Results

### 1. Species identification based on ITS analysis

ITS1 sequences were successfully obtained from 59 above-ground individuals and 163 soil seedbank-derived individuals (Table 1). All above-ground individuals were identified as *P. freyniana*. In contrast, soil seedbank-derived seedlings included not only *P. freyniana* but also *P. fragarioides* and *Potentilla cryptotaeniae* (Table 1, Fig. 2).

**Fig. 2.**
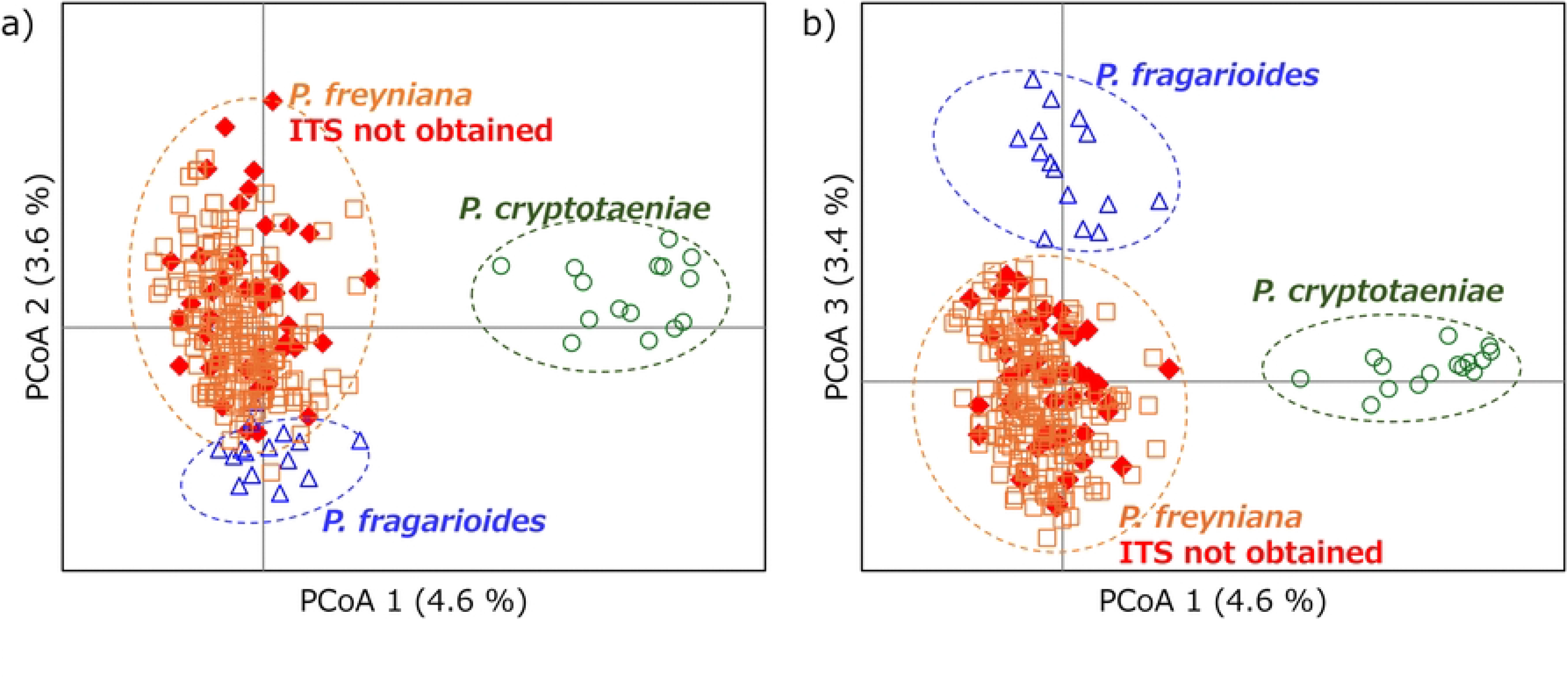
Principal coordinates analysis (PCoA) of 11 SSR genotypes used for species assignment of *Potentilla* individuals collected from above-ground vegetation and the soil seedbank. Orange squares indicate *P. freyniana*, blue triangles indicate *P. fragarioides*, and green circles indicate *P. cryptotaeniae*. Red diamonds represent individuals for which ITS sequences could not be obtained.

For individuals from which ITS1 sequences could not be obtained due to PCR amplification failure or other technical reasons, species identification was inferred using PCoA based on SSR data. All of these individuals clustered with samples identified as *P. freyniana* based on ITS1 sequences and were therefore treated as *P. freyniana* in subsequent analyses (Fig. 2).

### 2. Genetic diversity of above-ground and soil seedbank populations

*H*_E_ and *A*_R_ calculated from 15 SSR loci are shown in Fig. 3 and Table S2. In burned grasslands, both above-ground and soil seedbank populations exhibited high levels of genetic diversity (*H*_E_ = 0.831–0.864, *A*_R_ = 7.4–8.0). Although no above-ground individuals were detected in evergreen plantations, soil seedbank populations maintained similarly high genetic diversity (*H*_E_ = 0.831–0.853, *A*_R_ = 7.5–7.8). In contrast, the only above-ground population detected in deciduous plantations showed lower genetic diversity (*H*_E_ = 0.773,*A*_R_ = 6.5), whereas the corresponding soil seedbank populations retained higher diversity (*H*_E_ = 0.844–0.862, *A*_R_ = 7.8–8.2).

**Fig. 3.**
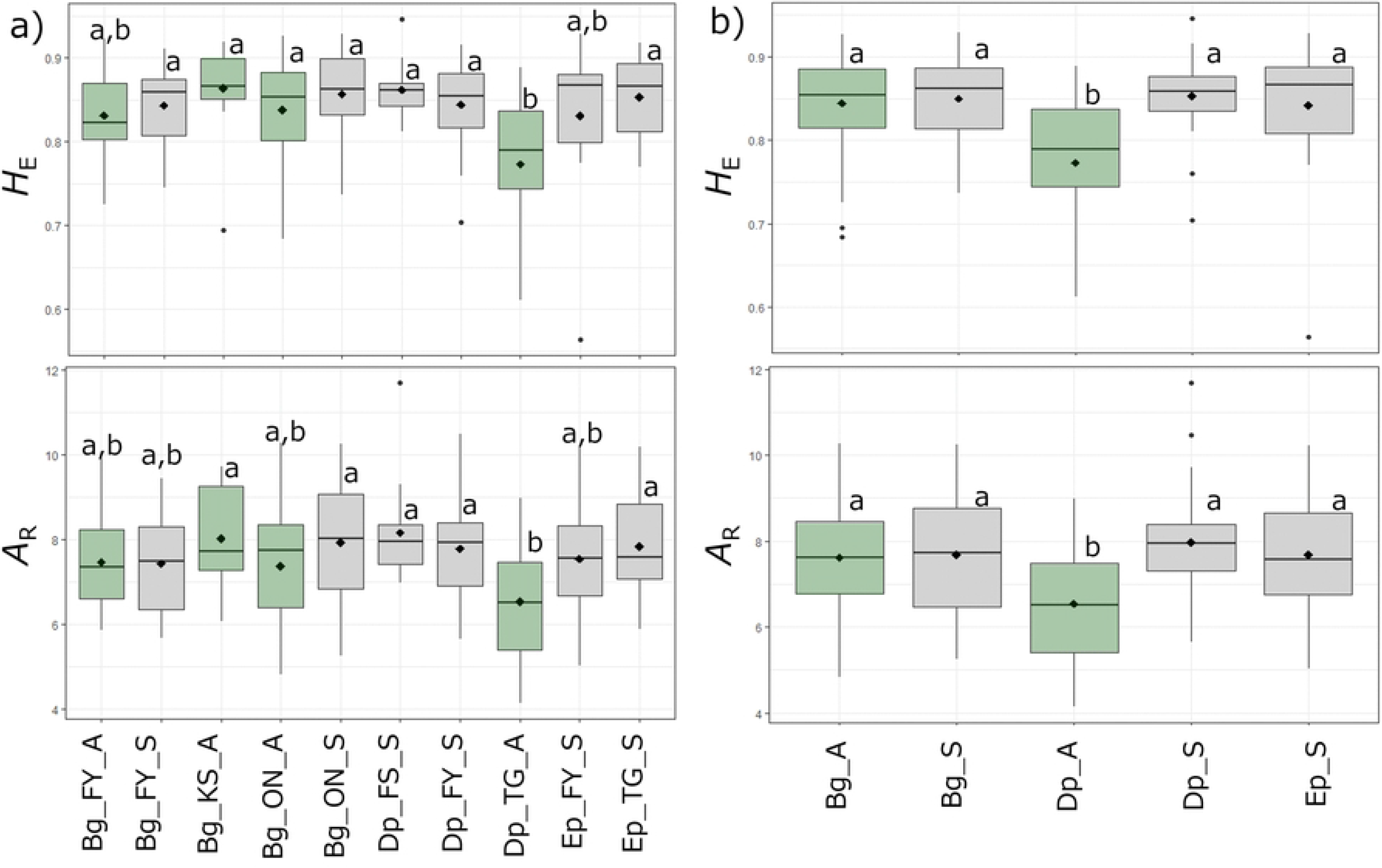
Distribution of expected heterozygosity (*H*_E_) and allelic richness (A_R_) among populations (a) and across land-use type and sample type (above-ground vs. seed bank) (b). Boxplots show locus- level values; boxes indicate the interquartile range, with the median shown as a central line. Whiskers extend to values within 1.5 times the interquartile range, and outliers are shown as open circles. Black diamonds indicate mean values. Different letters indicate significant differences among groups (Tukey’s HSD test, P < 0.05). Population codes represent land-use type, site, and sample origin. Land-use types are abbreviated as follows: Bg = burned grassland, Dp = deciduous plantation forest, and Ep = evergreen plantation forest. Site abbreviations (e.g., FY, KS, ON, FS, TG) indicate sampling locations. Sample origin is indicated by suffixes: A = above-ground individuals; S = seedlings derived from the seedbank. Box colors indicate sample origin (light green = above-ground populations; gray = soil seedbank populations).

Comparisons among categories revealed that the above-ground population in deciduous plantations (Dp_A) had significantly lower *H*_E_ and *A*_R_ than the other categories (Fig. 3b). In contrast, no significant differences were detected between soil seedbank populations and above-ground populations in burned grasslands.

Comparisons of PAR showed that, at the FY site, soil seedbank populations from burned grassland, deciduous plantation, and evergreen plantation exhibited similar values (Fig. 4). No substantial differences were detected between above-ground and soil seedbank populations, suggesting that private alleles were largely maintained within the soil seedbank. In contrast, at the ON burned-grassland site, PAR was significantly higher in the soil seedbank population than in the corresponding above-ground population.

**Fig. 4.**
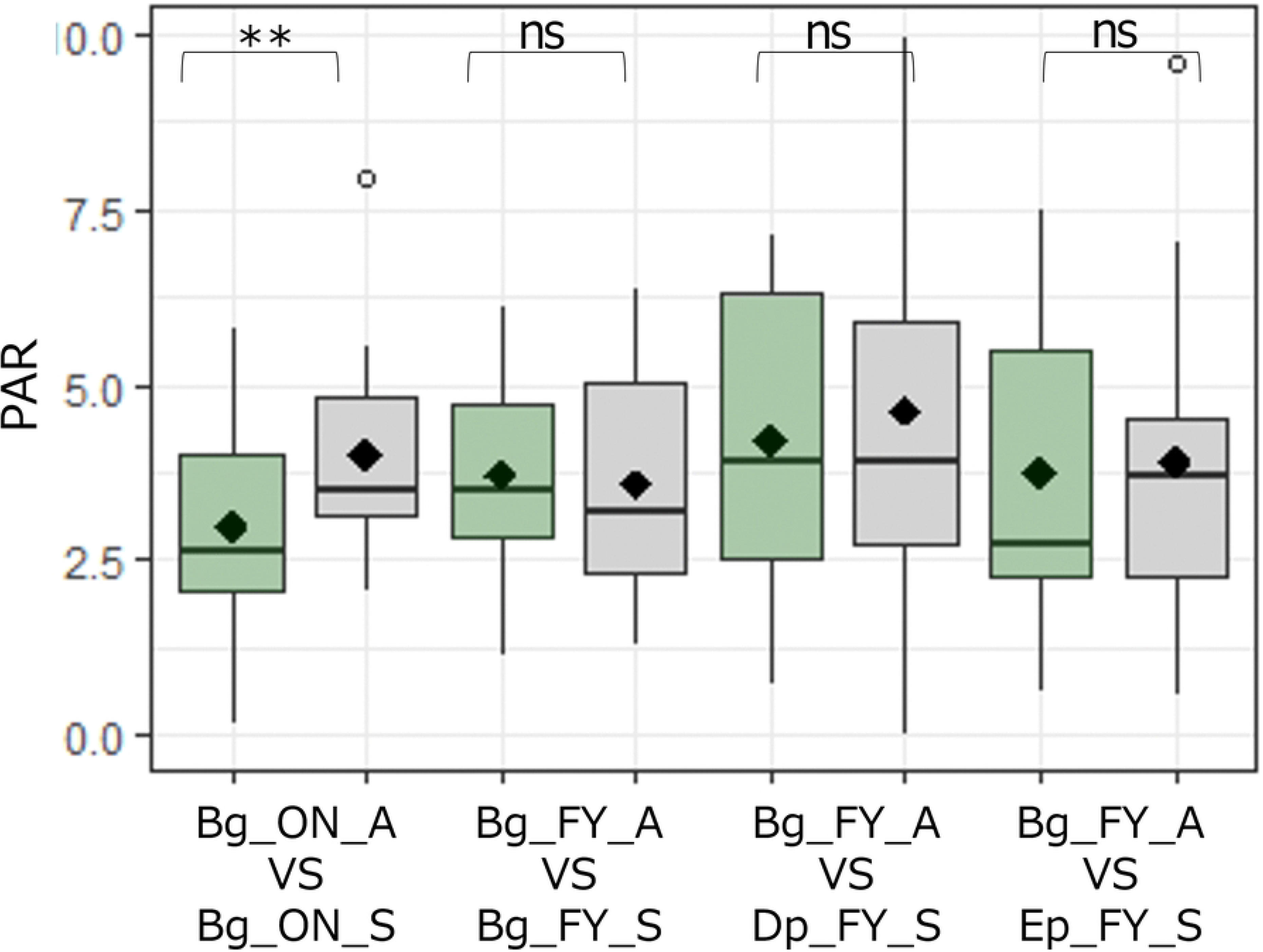
Comparison of private allelic richness (PAR) between above-ground and seed-bank samples within same sites, including grassland and plantation habitats. Land-use types are abbreviated as follows: Bg = burned grassland, Dp = deciduous plantation forest, and Ep = evergreen plantation forest. Site abbreviations (e.g., ON and FY) indicate sampling locations. Sample origin is indicated by suffixes: A = above-ground individuals; S = seedlings derived from the seedbank. Box colors indicate sample origin (light green = above-ground populations; gray = soil seedbank populations).

### 3. Population structure and genetic differentiation

Pairwise *F*_ST_ analyses revealed generally low levels of genetic differentiation among populations within localities, with *F*_ST_ values ranging from 0.004 to 0.072 (Table S3). No marked genetic differentiation corresponding to differences in land-use type was detected. Similarly, low *F*_ST_ values were observed between above-ground and soil seedbank populations within the same locality, indicating little genetic differentiation associated with land-use type.

AMOVA showed no significant genetic differentiation between above-ground and soil seedbank populations (*F*_RT_ = −0.003, P = 1.000; Table 3). In contrast, significant genetic differentiation was detected among populations (*F*_SR_ = 0.034, P = 0.001), although this component accounted for only 3.4 % of the total genetic variation. Most genetic variation was distributed within individuals (69.3 %) and among individuals within populations (27.3 %). These results suggest that above-ground and soil seedbank populations share largely the same gene pool.

**Table 3.** Analysis of molecular variance (AMOVA) showing genetic variation among and within above-ground and seed-bank populations of *Potentilla freyniana.* *F*_RT_ represents genetic differentiation between above-ground and soil seedbank groups, *F*_SR_ represents genetic differentiation among populations within groups, *F*_IS_ represents the inbreeding coefficient of individuals relative to their population, *F*_IT_ represents the inbreeding coefficient of individuals relative to the total dataset, and *F*_ST_ represents overall genetic differentiation.

| Source | df | Variance<br>component | Variation<br>(%) | Fixation index<br>(F-statistics) | *P* |
| --- | --- | --- | --- | --- | --- |
| Among groups (above-ground vs. seed bank) | 1 | 0.00 | 0.00 | $F_{RT} = -0.003$ | 1.000 |
| Among populations | 8 | 0.17 | 3.42 | $F_{SR} = 0.034$ | 0.001 |
| Among individuals | 208 | 1.36 | 27.29 | $F_{IS} = 0.283$ | 0.001 |
| Within individuals | 218 | 3.45 | 69.29 | $F_{IT} = 0.305$ | 0.001 |
| Total | 435 | 4.98 | 100.00 | $F_{ST} = 0.031$ | 0.001 |

In the STRUCTURE analysis, the Δ *K* statistic based on the method of Evanno et al. [38] showed a maximum value at *K* = 2, although a relatively high value was also observed at *K* = 7 (Fig. S1a). The log-likelihood [LnP(K)] increased with *K*, particularly up to approximately *K* = 4, after which it began to plateau (Fig. S1b). Therefore, multiple values of *K* were considered when interpreting population structure, and the results for *K* = 2–7 are presented (Fig. S1c). Based on individual cluster membership coefficients, above- ground individuals and soil seedbank-derived individuals from the same population generally showed similar patterns of cluster assignment, indicating concordant genetic structure between the two population types (Fig. S1c). However, admixture among multiple genetic clusters was observed in some individuals under all *K* values examined.

Shared-allele similarity estimated using Jaccard coefficients was generally high, and substantial differences among populations were not detected (Fig. 5). The highest levels of shared-allele similarity were not necessarily observed between above-ground and soil seedbank populations from the same site; similar levels of similarity were also found among populations from different sites. These results indicate that alleles were widely shared among populations and that there was no clear tendency for above-ground and soil seedbank populations within a site to exhibit exceptionally high similarity in allele composition.

**Fig. 5.**
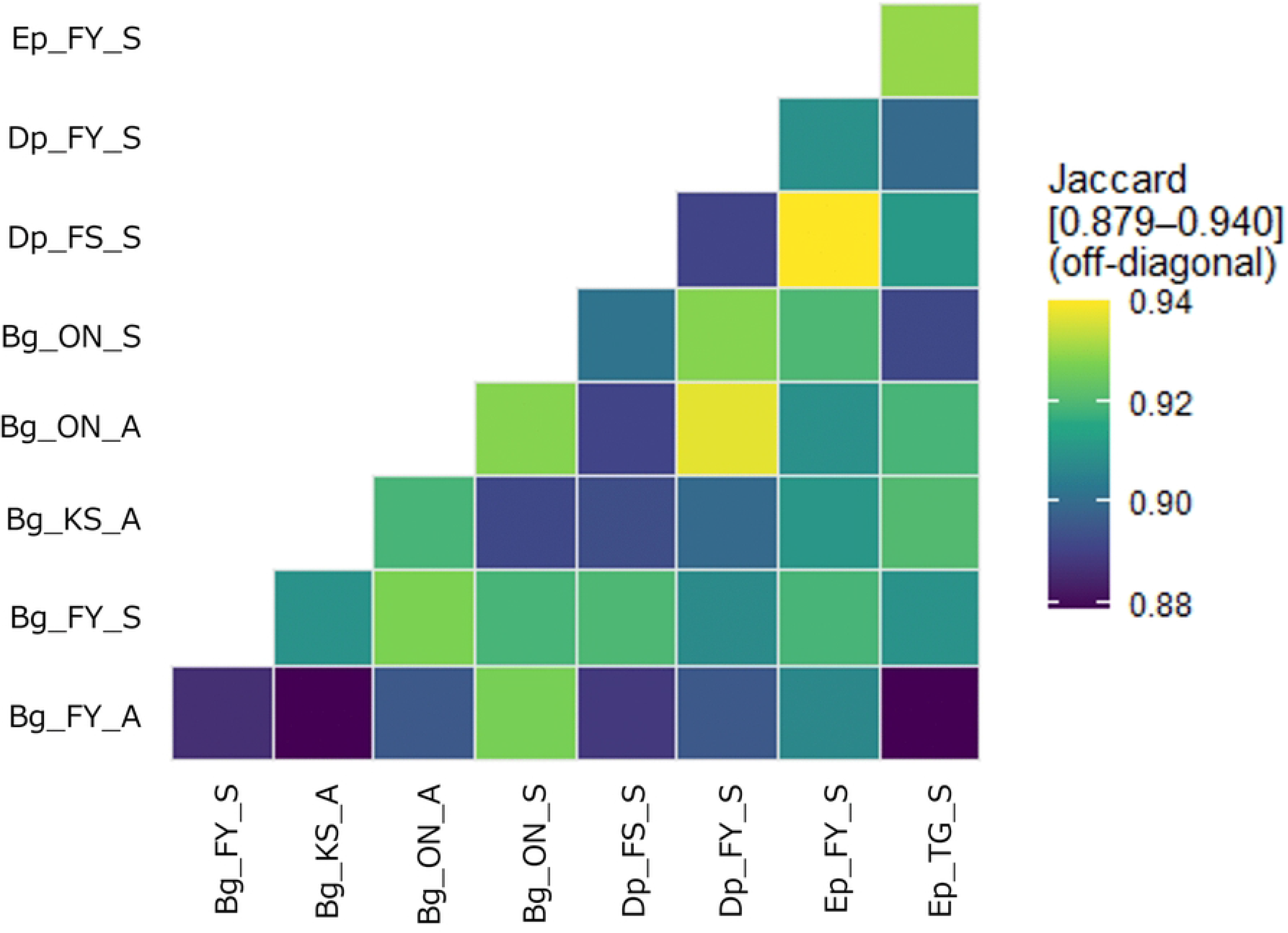
Heatmap showing pairwise shared allele similarity among populations, calculated using Jaccard coefficients based on presence/absence data. Population codes represent land-use type, site, and sample origin. Land-use types are abbreviated as follows: Bg = burned grassland, Dp = deciduous plantation forest, and Ep = evergreen plantation forest. Site abbreviations (e.g., FY, KS, ON, FS, TG) indicate sampling locations. Sample origin is indicated by suffixes: A = above-ground individuals; S = seedlings derived from the seed ban

## Discussion

### 1. Soil seedbanks persisting beneath plantation forests retain high levels of genetic diversity

The present study demonstrated that soil seedbank populations of *P. freyniana* retain high levels of genetic diversity even in areas where semi-natural grasslands have been converted to plantation forests. In particular, although no above-ground individuals were detected in evergreen plantation forests, the soil seedbank populations exhibited *H*_E_ and *A*_R_ values comparable to those observed in burned grasslands (Fig. 3). These findings indicate that genetically diverse seed populations can persist in the soil for several decades following conversion of semi-natural grasslands to plantation forests. The persistence of high genetic diversity in soil seedbanks within plantation forests aged approximately 21-62 years suggests that genetic variation derived from historical grassland populations can remain as a long-term genetic legacy. Although recent seed input from surrounding grasslands cannot be completely excluded, the analyzed seedlings originated from soil samples, including seeds recovered from the soil lower at 5–10 cm depth. Therefore, the high genetic diversity observed in plantation forests is consistent with the long-term persistence of soil seedbanks originating from historical grassland populations. Previous studies have shown that the effects of historical grasslands can persist in present-day vegetation following afforestation [11]. Our results extend this concept to the genetic level, suggesting that the genetic signature of historical grassland populations can persist through soil seedbanks.

In deciduous plantation forests, above-ground populations tended to exhibit reduced genetic diversity, whereas relatively high levels of diversity were maintained in the corresponding soil seedbank populations. This result suggests that assessments based solely on extant above-ground populations may underestimate the genetic resources that remain hidden within a region. Previous studies have shown that the legacy effects of historical grasslands can persist in contemporary species diversity and vegetation structure following land-use change [11]. Our results extend this concept to the genetic level, demonstrating that genetic diversity can also exhibit long-lasting legacy effects.

Soil seedbanks have been proposed to buffer the effects of genetic drift and population decline, thereby contributing to the maintenance of genetic diversity within populations [2]. The patterns observed in this study are consistent with such a mechanism. Therefore, even in former grasslands that have been converted to plantation forests, soil seedbank populations appear to function as important reservoirs of historical genetic diversity, especially for grassland species that form long-term persistent seedbanks. In addition, *P. freyniana* has been identified as a potential indicator species of species-rich semi-natural grasslands [40]. Therefore, the persistence of genetically diverse soil seedbank populations of this species may have implications not only for the conservation of *P. freyniana* itself but also for the restoration and conservation of species-rich grassland ecosystems.

### 2. Genetic continuity and differences between above-ground and soil seedbank populations

The results of this study suggest substantial genetic continuity between above-ground and soil seedbank populations. AMOVA revealed no significant genetic differentiation between above-ground and soil seedbank populations, indicating that they largely share the same gene pool (Table 3). Pairwise *F*_ST_ values were also low, further supporting weak genetic differentiation between the two population types. Similarly, STRUCTURE analyses showed no clear separation between above-ground and soil seedbank populations, and both exhibited comparable genetic compositions (Fig. S1).

In contrast, the shared-allele analysis revealed that, although a considerable proportion of alleles was shared between above-ground and soil seedbank populations, the degree of similarity varied among populations (Fig. 5). This finding suggests that soil seedbank populations do not simply represent the current above-ground populations, but rather integrate genetic information accumulated over multiple generations. Previous studies have shown that soil seedbanks can function as a genetic memory, maintaining levels of genetic diversity similar to those of above-ground populations while differing in allele frequencies [4].

Furthermore, at the ON burned-grassland site, private allelic richness (PAR) was significantly higher in the soil seedbank population than in the corresponding above-ground population (Fig. 4). This result suggests that soil seedbanks can retain alleles that have already been lost from contemporary above-ground populations or have become rare in them. Such differences may reflect the accumulation of genetic variation through the formation of soil seedbanks across multiple generations, as well as the effects of recent population decline and genetic drift in above-ground populations [1, 2].

These results indicate that soil seedbank populations do not constitute a gene pool independent of contemporary above-ground populations. Rather, they share a substantial proportion of genetic variation with current populations. Therefore, soil seedbanks appear to play an important role in linking past and present genetic diversity within grassland plant populations.

### 3. Implications of soil seedbanks for grassland restoration

The persistence of genetically diverse soil seedbank populations in plantation forests suggests that grassland restoration may be possible through the utilization of soil seedbanks in combination with management practices such as canopy opening, improvement of light conditions, and prescribed burning.

However, caution is needed when generalizing these results to all grassland species, as *P. freyniana* is a relatively common species that is widely distributed in semi-natural grasslands. *P. freyniana* is also recognized as a species that forms a long-term persistent seed bank, whereas many other typical grassland species in Japan are unlikely to establish such persistent soil seedbanks [20, 21]. In species that do not form persistent soil seedbanks, the seedbank is likely to disappear following afforestation as above-ground populations decline or disappear. Previous studies have reported that in rare or declining species, soil seedbank populations may harbor higher levels of genetic diversity than extant above-ground populations [15, 17]. In contrast, our results are consistent with those of Iberl et al. [41], who found similar levels of genetic diversity in above-ground and soil seedbank populations of a common species. One possible explanation is that common species are less susceptible to genetic erosion and population bottlenecks than rare or declining species.

A broader review by Honnay et al. [18] suggested that persistent soil seedbanks may mitigate the effects of habitat fragmentation by buffering populations against genetic drift and population differentiation, even when they do not contain substantially higher levels of genetic diversity than above-ground populations. The results of this study are consistent with this view. Genetic diversity in the soil seedbank was generally similar to that of above-ground populations. In addition, AMOVA and STRUCTURE analyses revealed little evidence of genetic differentiation between the two population types, indicating that soil seedbanks contribute to the long-term maintenance of genetic variation within populations.

These findings suggest that the conservation value of soil seedbanks may depend on species rarity, population dynamics, and life-history characteristics. For rare or threatened species, soil seedbanks may preserve genetic variants that have been lost or become rare in above-ground populations. For common species such as *P. freyniana*, soil seedbanks may instead play an important role in maintaining genetic diversity through time and providing a source of genetic resources for population recovery.

### 4. Conclusion

This study provides empirical evidence that genetically diverse soil seedbank populations can persist for several decades after the conversion of semi-natural grasslands to plantation forests. By integrating analyses of both above-ground and soil seedbank populations, we showed that soil seedbanks retain substantial genetic diversity and may contribute to the long-term conservation of genetic resources within grassland plant populations. However, our conclusions are based on a single grassland species that forms a long-term persistent seedbank, and the extent to which similar patterns occur in other grassland species remains unclear. Future studies should compare species with contrasting life-history traits and seedbank persistence, as well as examine whether the genetic diversity preserved in soil seedbanks contributes to successful population recovery following grassland restoration. Such studies will further clarify the role of soil seedbanks in conserving biodiversity and restoring grassland ecosystems under ongoing land-use change.

## Acknowledgements

We sincerely thank Issei Nishimura, Hiroki Yuasa, Yuki Iwachido, and James R. P. Worth for their assistance with field sampling. We are grateful to Chisako Furusawa and Miyabi Sato for their assistance with genetic analyses. We also thank the landowners for permitting access to their properties and for supporting our field investigations. This work was supported by JSPS KAKENHI Grant Number JP23K25051. The authors used Microsoft Copilot to assist with translation and English language editing of the manuscript. The authors critically reviewed, edited, and approved all generated content and took full responsibility for the accuracy and integrity of the manuscript.

## Supporting Information Captions

Fig. S1 Genetic structure of populations inferred by STRUCTURE. (a) Δ*K* values and (b) mean log-likelihood [LnP(K)] for different numbers of clusters (*K*). (c) Assignment probabilities of individuals to genetic clusters for *K* = 2–7, including both above-ground and seed-bank samples of *Potentilla freyniana* from the Kaida Plateau.

Table S1. Aligned ITS1 sequences (partial) showing nucleotide positions 114 and 127 of *Potentilla freyniana* and *P. fragarioides* from multiple geographic locations, including sequences obtained in this study and from GenBank.

Table S2. Genetic diversity indices (*N*_A_*, N*_E_*, I, H*_O_*, H*_E_*, uH*_E_*, F*_IS_, and *A*_R_) across above-ground and seed-bank populations of *Potentilla freyniana* based on 15 SSR loci.

Table S3. Pairwise *F*_ST_ values (lower triangle) and their statistical significance (upper triangle) among populations based on 11 SSR loci.

